# Using bobcat (*Lynx rufus*) movement patterns to inform strategies for road mitigation

**DOI:** 10.64898/2026.01.23.701409

**Authors:** Margaret Mercer, Cheryl Mollohan, Kerry Baldwin, Al LeCount, Michael J. Noonan, Jesse Alston

## Abstract

1. Even for wildlife species that often persist in urban and suburban areas, vehicle collisions remain a common source of mortality, and roads can fragment and degrade habitat. Quantifying animal behavior near roads can help wildlife managers develop management strategies to reduce mortality from vehicles while maintaining connectivity.
2. To determine how roads affect movement of bobcats (*Lynx rufus*)—a common mesopredator in urban and suburban areas of North America—we analyzed GPS tracking data from bobcats using continuous-time movement analyses. Our study focused on three questions regarding bobcat movement near roads: (1) Are roads barriers to bobcat movement? (2) How often do bobcats use wildlife crossing structures to cross roads? (3) How does bobcat movement behavior change when bobcats are closer to roads?
3. We found that bobcats crossed roads 11% less frequently than expected from random chance, and this effect was largely driven by interstates and major local roads. We found little evidence that bobcats selectively used culverts or underpasses to cross roads, or that bobcat movement behavior (i.e., speed or home range size) varied with road density, although daily distance traveled increased with road density.
4. *Synthesis and applications:* Managers attempting to reduce bobcat mortality from vehicle collisions must do more than simply building wildlife crossing structures. Fences to funnel bobcats toward crossing structures, rumble strips to scare bobcats from roads, reduced speed limits, and wildlife warning signs for drivers may be more effective tools for reducing bobcat-vehicle collisions. This study also provides a rigorous framework for considering the implications of movement behavior for lack of connectivity and mortality as distinct but not mutually exclusive threats posed by roads for wildlife.

## INTRODUCTION

Roads can act as barriers that limit animals’ ability to find food and mates (Broekman et al., 2024; Forman & Alexander, 1998; González-Bernardo et al., 2023; Ochs et al., 2024). This can reduce a population’s fitness (via genetic isolation and inbreeding) and capacity to adapt to changing environments (I. Keller & Largiadèr, 2003; L. F. Keller & Waller, 2002; Legrand et al., 2017). Even when animals can cross roads, they are at risk of being injured or killed in vehicle collisions, which have been shown to cause substantial mortality in birds (Grilo et al., 2020; Silva et al., 2020), reptiles (Jochimsen et al., 2014), amphibians (Ochs et al., 2024), and mammals (Silva et al., 2020) throughout the world (Grilo et al., 2021; Hill et al., 2019). Because of the dual threats of reduced connectivity and direct mortality, road ecology is an emerging discipline within conservation biology (Ament et al., 2023; Ford et al., 2022; Liczner et al., 2024; Tucker et al., 2023).

Conservation biologists who want to help animals cross roads and avoid vehicle collisions typically must change driver behavior, animal behavior, or both. Common methods of changing driver behavior include decreasing speed limits and posting wildlife warning signs for drivers (Lester, 2015). Wildlife behavior can be changed by installing fencing alongside roads, building wildlife crossing structures like overpasses and underpasses, and clearing roadside vegetation to make spaces near roads less appealing for wildlife (Lester, 2015; Rytwinski et al., 2016; Zheng et al., 2024). Installing rumble strips can have the dual effect of making drivers slow down and warning wildlife that a car is nearby (Lester, 2015). To understand the extent to which each of these mitigation measures may be effective, it is important to study how wildlife interact with mitigation measures and roads in general.

Specific behaviors that wildlife exhibit in response to roads can inform management actions (Gagnon et al., 2011; Jacobson et al., 2016; Matos et al., 2019). For example, if a road is acting as a barrier to animal movement, managers can focus on facilitating connectivity. Conversely, frequent road crossings would suggests reducing road mortality risk is a greater concern. If animals actively search for and use existing crossing structures to cross roads, then installing more of these structures might be effective for increasing connectivity. If animals crossing roads do not often use existing crossing structures, however, then focusing on reducing mortality by lowering speed limits and installing rumble strips may be more effective. Differences in cost and feasibility of mitigation strategies also make research into animal behavior necessary. Some road mortality mitigation measures are more expensive than others in terms of absolute costs (e.g., crossing structures are more expensive than fencing, which is more expensive than signage; Dumalakas et al., 2025; Gagnon et al., 2015; Huijser et al., 2009), so to understand what measures we should be implementing, we need to understand the primary risks wildlife are experiencing.

Like many mesocarnivores (Recio et al., 2015; Rodriguez et al., 2021), bobcats (*Lynx rufus*) often inhabit urban areas (Young et al., 2019) but are threatened by roads (Barthelmess & Brooks, 2010), which can reduce fitness via both reduced habitat connectivity (Crooks, 2002) and direct mortality (Dyck et al., 2023). Bobcats that avoid roads experience reduced habitat connectivity (Crooks, 2002; Litvaitis et al., 2015; Mayer et al., 2022), leading to lack of connectivity between populations, reducing gene flow (Ceia-Hasse et al., 2018; Lee et al., 2012), and therefore making populations less stable (Crooks, 2002). Bobcat-vehicle collisions are often a leading cause of mortality for bobcats (Bencin et al., 2019; Chamberlain et al., 1999; Fuller et al., 1995; L. R. Jones et al., 2020; Litvaitis et al., 1987), both in rural (Nielsen & Woolf, 2002) and urban (Serieys et al., 2021) areas. Studying movement patterns of urban bobcats around roads can therefore help us understand how to mitigate impacts of roads to improve bobcat conservation.

We leveraged a dataset of 35 GPS-tracked bobcats (*Lynx rufus*) and used continuous-time movement models to assess the influence of roads on bobcat movement behavior in a large city in the southwestern United States and inform potential conservation actions to mitigate the influence of roads on bobcats. To do this, we centered our study on three questions regarding bobcat movement around roads: (1) Are roads barriers to bobcat movement? (2) How often do bobcats use wildlife crossing structures to cross roads? (3) How does bobcat movement behavior change when bobcats are closer to roads?

## METHODS

### Study Area

The study area encompassed much of the western portion of the Tucson metropolitan area (which lies on the traditional homelands of the Tohono O’odham people) in Arizona, USA, bounded to the east by Interstates 10 and 19 (Fig. 2). Rainfall in the Tucson area is 27 cm/year (US Department of Commerce, 2024), with a mean high temperature of 38.4° C in June and a mean low temperature of 4.7° C in December (US Department of Commerce, 2025). Tucson experiences bimodal annual rainfall, with summer and winter rains and a hot, dry season in early summer. This area is characterized by Upland Sonoran Desert vegetation, such as saguaro (*Carnegiea gigantea*), fishhook barrel cactus (*Ferocactus wislizenii*), creosote bush (*Larrea tridentata*), foothills palo verde (*Parkinsonia microphylla*), and velvet mesquite (*Neltuma velutina*). Our study area covers a gradient of human disturbance, from relatively undisturbed open space and parks (e.g., Saguaro National Park and Tucson Mountain Park) to residential and industrial areas. The Santa Cruz River runs along the eastern border of our study area. Stretches of the Santa Cruz flow year-round due to releases of treated wastewater, but much of it is dry most of the year. Tributary washes flow intermittently and frequently serve as wildlife corridors through Tucson and can be useful passages for wildlife to navigate urban areas (Burnett, 2025).

**Figure 1:**
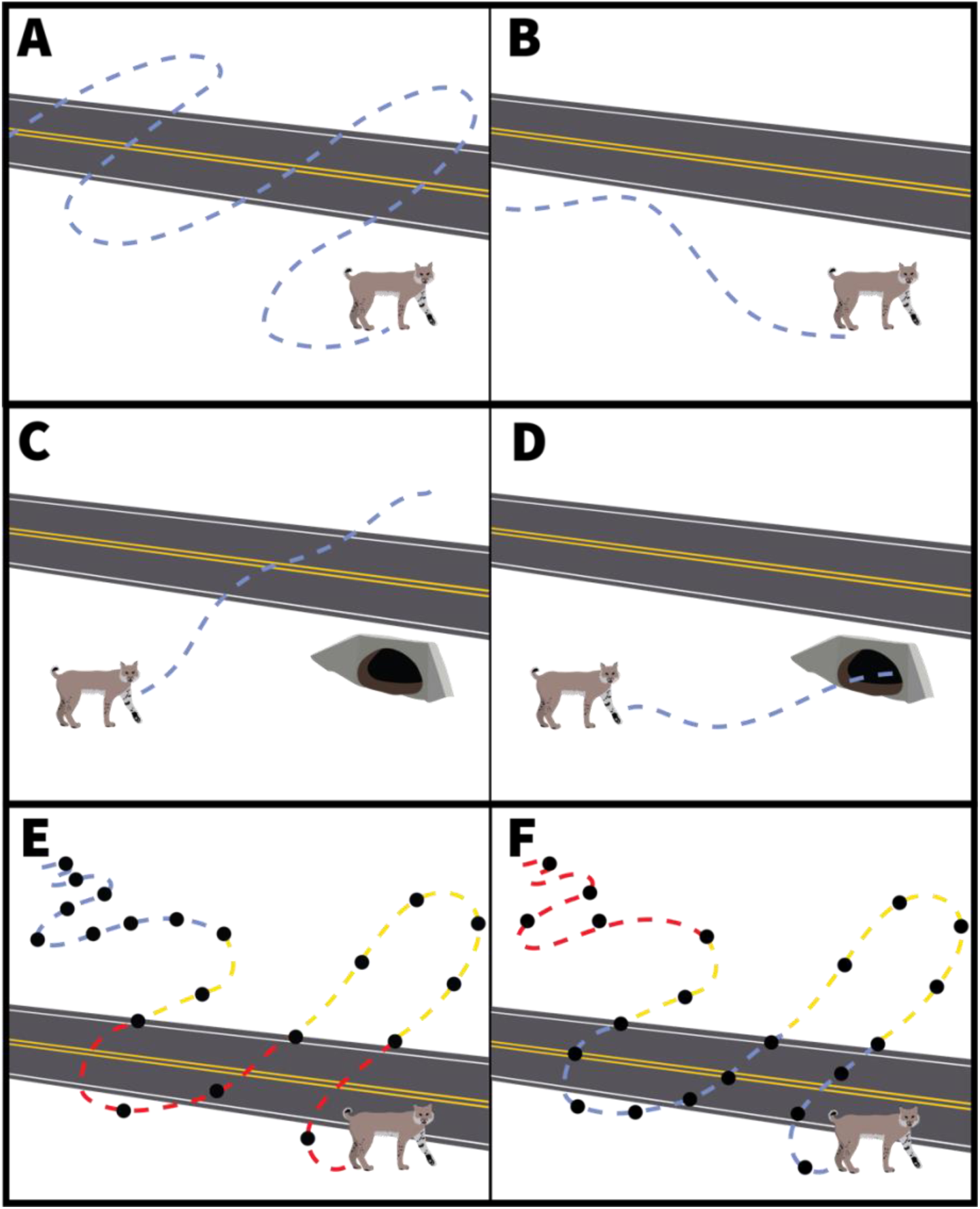
Schematic diagram showing possible bobcat responses to roads. Bobcats may either cross roads frequently (A) or avoid roads (B). Crossing roads frequently would imply that managers should focus on reducing mortality, while avoiding roads would imply that managers should focus on maintaining connectivity. Bobcats may cross roads upon encountering roads (C) or seek out crossing structures (e.g., culverts; D). If they do not seek out crossing structures, managers should focus on other mitigation strategies. If they seek out crossing structures, managers should install at least one crossing structure per bobcat territory. Bobcats may change their behavior around roads, such as by speeding up (E) or slowing down (F) when they approach roads.

**Figure 2:**
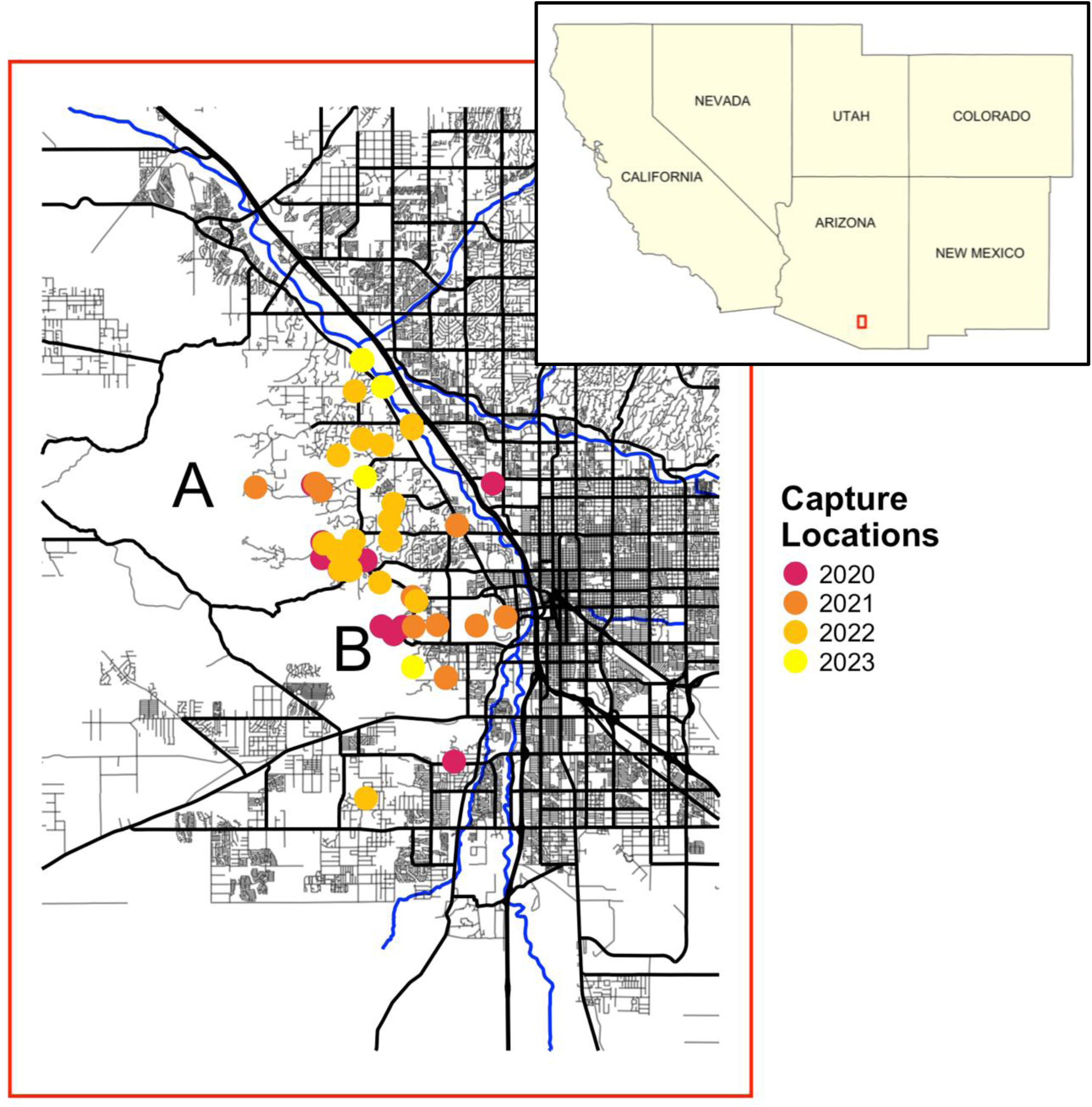
Map of study area in Tucson, Arizona, USA. Bobcat capture locations are colored by year. Blue lines represent rivers. Bolder black lines represent major roads (interstate, state, frontage, and major local roads), and thinner gray lines represent minor roads (minor local roads). Blue lines represent main riverbeds in the study area. Large roadless spaces are protected areas such as Saguaro National Park (A) and Tucson Mountain Park (B).

### Animal Capture

We conducted capture operations from November to April from 2020 to 2023. We captured bobcats using cage traps baited with roadkill, sight attractants, and commercial lures. All capture and handling protocols followed the recommendations of the American Society of Mammalogists’ guidelines for the use of wild mammals in research (Sikes & the Animal Care and Use Committee of the American Society of Mammalogists, 2016).

A veterinarian immobilized bobcats with ketamine (4 mg kg^-1^ body weight), Dexdomitor (0.04 mg kg^-1^ body weight), and Butorphanol (0.4 mg kg^-1^ body weight; Kreeger & Arnemo, 2018), placing a blindfold and protective optical lubricant over their eyes during immobilization. We recorded weight and sex and estimated age by tooth replacement and wear. For all bobcats weighing ≥ 5.5 kg (n = 35), we placed custom fitted programmable radio collars with GPS and VHF capability (Telonics TGW-4277 Iridium, Mesa, AZ, USA) and a breakaway mechanism (Telonics CR-7B, Mesa, AZ, USA) programmed to detach before the end of battery life. We fitted three bobcats weighing < 5.5 kg with smaller radio collars (Telonics TGW-4177 Iridium, Mesa, AZ, USA) and breakaway mechanisms (Telonics CR-5B, Mesa, AZ, USA). Radio collars weighed 280 and 140 g each, respectively. After collaring each bobcat, we reversed the effects of Dexdomitor using Atipamazole (0.2-0.4 mg kg^-1^ body weight; Kreeger & Arnemo, 2018) and monitored the bobcat until it was stable on its feet (1.5-2 hours).

Between November 2020 and December 2023, we captured 56 different bobcats 68 times: 21 adult males, 25 adult females, 2 subadult males, and 8 bobcats of unknown sex (1 subadult, 7 kittens). Thirty-eight bobcats were radio-collared: 36 adults and 2 subadults, of which 14 were male and 24 were female. In total, we recorded 53,683 bobcat locations across these individuals. The length of time radio-collars were deployed on bobcats ranged from 1 day to 2.5 years, with mean and median values of 1 year. The sampling interval of GPS locations ranged from 2 to 13 hours, with an average of 5.4 hours and a median of 4 hours.

### Statistical Analysis

We assessed GPS locations and excluded those exhibiting a dilution of precision > 6 (Van Sickle, 2015) and filtered out locations we deemed implausible based on travel distance between locations. Three bobcats with < 50 GPS fixes were excluded from analysis.

To map roads, we used road data from the Pima County Geospatial Data Portal (*Pima County Open Data*, 2024). We distinguished between major roads (i.e., interstate, state, frontage, and major local roads) and minor roads (i.e., minor local roads) in our analyses. We located crossing structures in our study area using the Arizona Department of Transportation “National Bridge Inventory Bridge and Culvert Conditions” dataset (*ADOT NBI Bridge and Culvert Conditions*, 2024). We visually assessed each crossing structure using Google satellite imagery to validate the location of crossing structures and to identify the type of crossing structure. We also added 17 unmarked crossing structures that we discovered during this process. Within the study area, there were 83 crossing structures, including places where roads passed over culverts (n = 34), washes (n = 38), and other roads (n = 11) (*ADOT NBI Bridge and Culvert Conditions*, 2024).

We used the ‘ctmm’ package (v1.2.0; Calabrese et al., 2016; Fleming & Calabrese, 2023) to conduct continuous-time movement modeling and calculate movement metrics. We first calibrated measurement error of the collars and removed outliers (Fleming et al., 2021). We selected a continuous-time movement model that best fit each individual’s movements using second-order Akaike’s Information Criterion (Burnham & Anderson, 2003; Calabrese et al., 2016). We used this model to calculate the average daily movement speed and predicted movement path (for each bobcat) and instantaneous movement speed (for each GPS location; Noonan et al., 2019, 2022). For each individual that was range-resident (33 out of 35 individuals), we estimated the home range using Autocorrelated Kernel Density Estimation (AKDE; Silva et al., 2022) to identify the 95% coverage area (Fleming et al., 2015). We also summed the total length of all roads within each individual’s home range (*Pima County Open Data*, 2024) and divided it by the home range’s area to estimate road density in each bobcat’s home range.

We identified broad patterns in bobcat movement patterns before answering our specific questions. We assessed the correlation between road density within a bobcat’s range and the number of times that bobcat crossed roads per day using a linear regression. We used t-tests to compare home-range size, number of road crossings per day, road density within the bobcat’s home range, frequency of crossing structure usage, and average movement speed between males and females.

We then simulated movement data for each bobcat to represent the null hypothesis that bobcats do not respond to roads. To simulate road-agnostic bobcat movement paths, we used the ‘simulate’ function in ctmm (Noonan et al., 2022), which, for each bobcat, matches the centroid of the home range, the selected movement model, and the amount of time the collar was deployed. We ran 1000 simulations for each bobcat.

#### (1) Are roads barriers to bobcat movement?

Using the GPS movement data and our map of roads in the Tucson area, we calculated the total number of times each bobcat’s most likely path crossed a road. We calculated the mean number of road crossings across all 1000 simulations for each bobcat. We compared the number of observed road crossings to the mean number of simulated crossings using a paired Wilcoxon signed-rank test to determine whether our observed bobcats crossed roads more or less frequently than would be expected if bobcats showed no response to roads. We performed this analysis for all roads, then for major and minor roads separately.

#### (2) How often do bobcats use wildlife crossing structures to cross roads?

We recorded the coordinates of both ends of each crossing structure within our study area and used the ‘sf’ package (v1.0.16; Pebesma, 2018) to connect the end coordinates to create a line representing each crossing structure. Using the ‘geosphere’ package (v1.5.18; Hijmans et al., 2024), we identified every crossing event on the road within 7 m, which reflects the median location error for the GPS radio-collars used in this study, of a crossing structure for both real and simulated bobcats (Noonan et al., 2022). We then compared the number of observed crossings near structures to the mean number of simulated crossings near structures (i.e., a null model of no preference toward crossing structures) using a paired t-test.

#### (3) How does bobcat movement behavior change when they are closer to roads?

For each GPS location, we evaluated the relationship between distance from roads and the instantaneous speeds of the bobcat (in m sec^-1^; Noonan et al., 2022) using a linear mixed model. We performed the analysis for major roads only, minor roads only, and both road types combined. We log-transformed distance from roads, because distance is much more likely to affect instantaneous speeds when bobcats are very close to roads than when they are very far from roads. We incorporated random slopes and intercepts for each bobcat to account for non-independence within individual bobcats and fit models using the ‘lme4’ package (v 1.1.35.5; Bates et al., 2025). We also evaluated the relationship between road density in each bobcat’s home range and its home range size and average daily distance traveled using linear models. We used the ‘lmerTest’ package (v 3.1.3; Kuznetsova et al., 2017) for these statistical tests.

All analyses were conducted in the R statistical software environment (v4.4.1; R Core Team, 2023).

## RESULTS

### (1) Are roads barriers to bobcat movement?

**Bobcats crossed roads 11.6% less frequently than expected under the null hypothesis that roads did not alter movement behavior.** On average, bobcats crossed roads 3,050 times (range: 46-8,274), while simulated bobcats with identical movement parameters crossed roads 3,452 times (Wilcoxon test; *V* = 207, *p* = 0.078; Fig. 3a). This avoidance behavior was predominantly driven by major roads—bobcats crossed major roads 26.5% less frequently (474 crossings, range: 0-2,020) than expected (645 crossings; Wilcoxon test; *V* = 114, *p* = 0.00063; Fig. 3b). They did not cross minor roads less frequently (2,576 crossings, range: 33-7,715) than expected (2,807 crossings; Wilcoxon test; *V =234*, *p* = 0.19; Fig. 3c).

**Figure 3:**
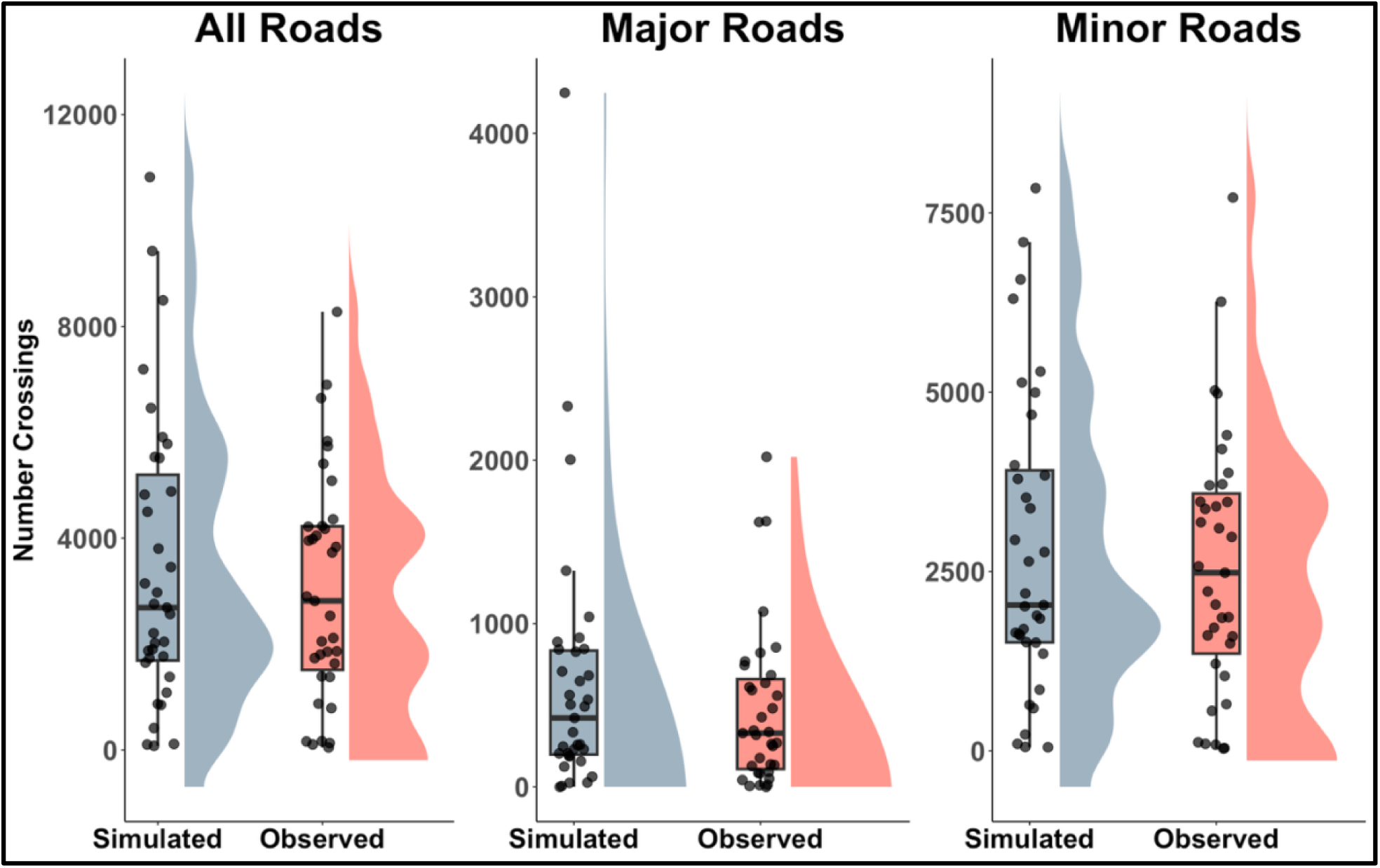
Frequency distributions of the total number of observed and simulated road crossings by bobcats in an urban area in Tucson, AZ, USA. Simulated crossings represent the null hypothesis of no road avoidance. Each point represents the total number of road crossings by individual. The middle horizonal line of each box-and-whisker plot indicates the median value for each group, the boxes indicate the central 50% of each group, and the whiskers indicate the lower and upper 25% of each group.

The number of road crossings by bobcats was positively related with road density within their home ranges (β = 0.68; 95% CI: 0.50–0.87; *p* < 0.001). Male home ranges were 2.5 times larger than female home ranges (mean = 17.1 km^2^ for males [range: 9.5-28.4 km^2^], 6.8 km^2^ for females [range: 2.1-15.6 km^2^]), but we found no evidence for differences between sexes for the number of road crossings per day (*t* = –0.49, df = 29.3, *p* = 0.63), road density within home ranges (*t* = –0.79, df = 28, *p* = 0.44), percentage of crossings near a crossing structure (*t* = –0.087, df = 28, *p* = 0.93), or daily distance traveled (*t* = 1.13, df = 16.6, *p* = 0.28).

### (2) How often do bobcats use wildlife crossing structures to cross roads?

**Bobcats did not use crossing structures to cross roads more than expected under the null hypothesis that roads did not alter movement behavior.** The mean distance from each point where a bobcat crossed a road to the nearest crossing structure was 1,804 meters (95% CI: 1,796-1,812), which was less than the 1,990 meters (95% CI: 1,975-2,005) for simulated bobcats (*t* = – 21.1, df = 55,173, *p* < 0.001), but the difference is not likely to be biologically meaningful. Only 1.3% (95% CI: 0.2%-2.4%) of observed bobcat road crossings occurred within 7 m of crossing structures compared to 0.4% (95% CI: 0.16%-0.6%) of simulated road crossings (paired t-test; *t* = –1.93, df = 34, *p* = 0.06). Two bobcats that lived in the Santa Cruz River corridor were outliers, using crossing structures at rates of 15.7% and 11.3% because they frequently crossed under bridges while traveling up– and downriver. When these two bobcats were excluded, the other bobcats used crossing structures 0.6% of the time (95% CI: 0.1%-1.0%) compared to 0.3% (95% CI: 0.1%-0.4%) of simulated road crossings (*t* = –1.68, df = 29, *p* = 0.10). Eleven of 35 bobcats (31.4%) never crossed a road within 7 m of a crossing structure.

### (3) How does bobcat movement behavior change when they are closer to roads?

**We did not find evidence that movement speed or home range size vary in response to roads.** Bobcat movement speed did not vary with proximity to roads in general (β = –0.00014, 95% CI = –0.0009 – 0.0007; *p* = 0.73; Fig. 4a) or minor roads (β = 0.00015, 95% CI = –0.0006 – 0.0009; p = 0.7), and though bobcats did slow down as they approached major roads (β = – 0.0015, 95% CI = –0.0023 – –0.00064; p = 0.0015), this reduction in speed not likely to be biologically meaningful. Similarly, home-range size was not related to road density (β = –0.013, 95% CI = –0.45 – 0.42; *p* = 0.96; Fig. 4b). Daily distance traveled, however, was positively related to road density within an individual’s home range (β = 0.25, 95% CI = 0.04 – 0.46; *p* = 0.028; Fig. 4c).

**Figure 4.**
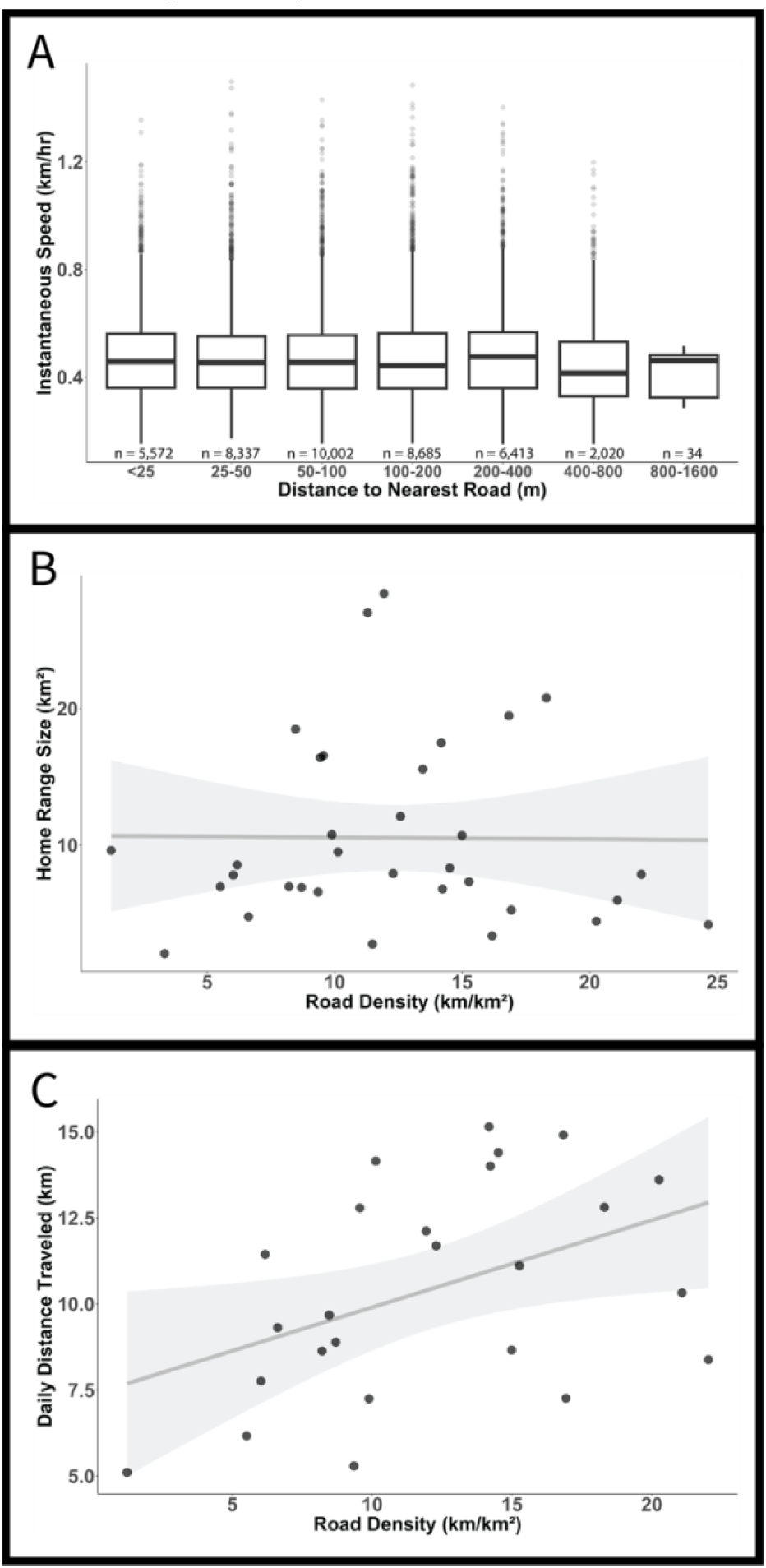
Relationships between bobcat movement metrics and roads in an urban area in Tucson, AZ, USA. Panel A shows that proximity to roads has no effect on instantaneous speeds of bobcats. The middle horizonal line of each box-and-whisker plot indicates the median value for each group, the boxes indicate the central 50% of each group, and the whiskers indicate the lower and upper 25% of each group. Points indicate outliers for each group. Panel B shows that there is no relationship between bobcat home range size and within-home range road density. Panel C demonstrates that the relationship between bobcat daily distance traveled and within-home range road density is positive. In panels B and C, points represent values for individual bobcats, dark gray lines and light gray ribbons represent linear relationships and 95% standard errors, respectively.

## DISCUSSION

Roads can impact animals by reducing connectivity and increasing mortality, and studying animal behavior can inform effective mitigation strategies for these conservation problems. Our analyses suggest that efforts by managers to increase connectivity for bobcats by building crossing structures are relatively unlikely to be effective, because bobcats did not seek out crossing structures, using them for only ca. 1% of road crossings, which was not significantly different from the expectation from random chance. Efforts by managers to reduce vehicle mortality by reducing speed limits, adding fencing, or installing warning signs or rumble strips are therefore more likely to be effective, particularly on minor roads like residential streets.

Bobcats crossed roads less frequently than expected by random chance, but this behavior was driven by major roads (i.e., interstate, state, frontage, and major local roads). This suggests that while major roads may reduce connectivity for bobcats, minor roads may be more likely to cause direct mortality from vehicle collisions (five of our thirty-eight study animals were killed in vehicle collisions during our study). That bobcats will readily cross minor roads corroborates the results of previous studies—animals overall avoid roads more when traffic increases (Collins et al., 2022; Hill et al., 2021; Tucker et al., 2023) and mesocarnivores in Central California (Serieys et al., 2021) and the Netherlands (van Langevelde et al., 2009) die more often on less-trafficked roads. Bobcat avoidance of major roads suggests that the cues used in avoiding major roads could be used to reduce vehicle mortality along minor roads (albeit at the expense of connectivity). This could include placing rumble strips on roads (Lester, 2015), installing alarms to go off when cars drive by (St. Clair et al., 2019), and clearing vegetation around roads (Lester, 2015). However, we found no evidence that bobcats changed their travel speed as they approached minor roads, and little evidence that they slow down as they approach major roads, which suggests that bobcats did not pause to assess road safety before approaching. This further indicates that changing driver behavior (e.g., reducing speed limits, posting wildlife warning signs, and adding speed bumps and chicanes to roads; D. N. Jones et al., 2014) may be more effective at mitigating bobcat mortality than deterring bobcats from crossing minor roads.

We found little evidence that bobcats selectively used crossing structures when crossing roads. Our study supports previous research showing that bobcats cross roads wherever they are encountered, sometimes despite the presence of crossing structures nearby (Hanley, 2022; Rytwinski et al., 2016; Tigas et al., 2002). Bobcats primarily cross roads in areas bordering preferred habitat (Cain et al., 2003), and in our study as well as others (e.g., Serieys et al., 2021), bobcats were most likely to use crossing structures when they fell along their path of travel, such as when a bobcat was traveling along a wash and crossed under a bridge. Even when animals use crossing structures, these structures do not always reduce the total number of vehicle mortalities (Elliott & Stapp, 2007). Pairing crossing structures with fencing increases the probability of animals using those structures (Rytwinski et al., 2016). Installing short (100-m) stretches of fencing on either side of crossing structures may increase crossing structure use by bobcats in areas where bobcats already use crossing structures, but fencing may have little effect on crossing structure usage in areas where bobcats do not already use them (Cain et al. 2003).

Managers can identify movement corridors to guide crossing structure placement and fencing at those locations (Cain et al., 2003), but this is not likely to be effective for bobcats. Bobcats are solitary and territorial and do not generally leave their territories (Bailey, 1974; Benson et al., 2004; Lariviere & Walton, 1997). Areas occupied by bobcats typically host two adults at a time: a female and a male whose territory overlaps multiple females (Bailey, 1974). Crossing structures are therefore only likely to help the few individuals in whose territories the crossing structure is placed, and maintaining meaningful connectivity would likely require installing crossing structures in multiple bobcat territories. Crossing structures may also benefit subadults dispersing from their natal ranges, but collecting data on natal dispersal to evaluate subadult bobcat use of such structures is challenging (Janečka et al., 2007). Future studies on natal dispersal may reveal the extent to which crossing structures can facilitate population-wide genetic connectivity across barriers like major roads.

Findings on how bobcats alter movement in response to changing road density are inconsistent: Riley et al. (2003) found that bobcat home-range size increased with urbanization, but both our study and Tigas et al. (2002) found no change in home-range size as road density or habitat fragmentation increased. Tigas et al. (2002) proposed that the reduction in habitat quality that occurs with urbanization could be offset by human-subsidized food sources such as trash or pets. In our study area, many people maintain native brushy vegetation, bird feeders, and water sources, which bobcats in our study were often observed visiting. This supports the human subsidy hypothesis and may explain the negligible relationship between bobcat home range size and road density. In contrast, bobcats in our study moved farther each day in areas with higher road density, perhaps due to increased vigilance in urban areas resulting from fear of human activity. Bobcats also tend to travel more quickly through habitat that is less favorable (Abouelezz et al., 2018; Dickson et al., 2005), indicating that higher densities of roads decreases habitat suitability. Similarly, urban mountain lions reduce time spent feeding when they hear humans (Smith et al., 2017), which increases the rate at which they must kill prey to survive. Together, this implies that rather than increasing the size of their territories, urban bobcats may move more within their territories to avoid humans and to seek more frequent opportunities to hunt. They may also need to compensate for the higher spatial variance in resource availability that occurs in urban environments, moving quickly through resource-poor areas (e.g., roads, houses, and parking lots) to get to higher-value patches (e.g., bird feeders and water sources). In addition, if territory sizes do not change with changing road densities but daily distance traveled does, urban bobcats traverse their home ranges more frequently than bobcats in areas of lower road densities, which in turn likely causes urban bobcats to cross roads even more frequently, offsetting their slight avoidance of roads and further corroborating that vehicle mortality poses a larger threat to urban bobcats than reduced connectivity from roads.

Overall, maintaining connectivity for bobcats in urban areas while minimizing vehicle mortality will be difficult: bobcats rarely avoided crossing roads and did not preferentially seek out crossing structures. Management efforts aimed at reducing mortality are therefore more likely to benefit bobcats than efforts intended to increase connectivity. More generally, this study demonstrates how managers can use studies of movement behavior to choose management strategies that are most likely to be successful. Similar methods can be applied to other species facing conservation challenges from roads and help managers mitigate the negative effects of roads on those species.

## Acknowledgements

This study was funded by an Arizona Game and Fish Department Heritage Program Urban Wildlife Grant (Lottery Dollars for Wildlife), private donations facilitated by the Southwest Wildlife Conservation Center, the School of Natural Resources and the Environment at the University of Arizona, the Martha I. Grinder Scholarship in Wildlife and Fisheries Resources, the Arrington Memorial Scholarship, the American Society of Mammalogists Grant-in-Aid of Research, and the University of Arizona Graduate Professional Student Council. Jeff Gagnon and Colin Beach contributed knowledge of road ecology of bobcats, Jeffrey Oliver and Ellen Bledsoe assisted with coding the analysis and data wrangling, and Javan Bauder and Ellen Bledsoe contributed ideas that greatly strengthened the manuscript.

## Author contributions

CM, KB, and AL designed the field study, collected the data, and provided a write-up of the methods. MM and JMA analyzed the data and led the writing of the manuscript. CM, MM, MJN, and JMA contributed to the drafts and gave final approval for publication.

The authors declare no conflict of interest.

